# Solitary phytoplankton cells sink in the mesopelagic ocean

**DOI:** 10.1101/2025.03.16.643536

**Authors:** Annie Bodel, Margaret Estapa, Colleen A. Durkin

## Abstract

Phytoplankton, and their carbon, are typically exported from the surface ocean when packaged inside larger, sinking detrital particles. This process draws carbon out of the atmosphere, where it can be sequestered for long time periods in the deep ocean. Phytoplankton can also sink as solitary cells, but direct observations are scarce and the ecological significance is unknown. We collected unprecedented observations of solitary sinking cells during month-long observations in the upper 500 m at two contrasting ocean locations. While these cells account for only a small fraction of the total particulate organic carbon flux (<5%), they provide essential nutrients and a persistent source of food to deep sea ecosystems while preserving a seed bank for future phytoplankton blooms. In one case, observed depth changes over time allowed calculation of a sinking speed of 6 m d^-1^. The disaggregation of detrital aggregates collected at the same time could not account for the magnitude or composition of individually sinking cells in our samples, although in some instances their fluxes were correlated. Instead, the data imply that these solitary cells were transferred through the upper mesopelagic as individually sinking particles and play key roles in ocean ecology and biogeochemistry.

## Introduction

Phytoplankton growth in the surface ocean generates organic carbon that feeds ocean ecosystems and initiates a pathway that sequesters carbon away from the atmosphere and into the deep ocean (1,2). Most of that sequestration occurs through the gravitational sinking of organic particles, including fecal pellets, detrital aggregates, and animal carcasses or cast-off parts. This collection of processes resulting in deep carbon sequestration is called the biological carbon pump and is both a key mechanism regulating global climate and the primary food supply for deep sea ecosystems. Our ability to accurately account for this system of processes guides how ocean carbon cycle models are created and how future ocean monitoring efforts are designed. If we overlook key processes of the biological carbon pump, our ability to respond and adapt to a changing climate will be limited. Phytoplankton production is a key driver of the biological carbon pump, and it is assumed that these microscopic cells contribute directly to downward fluxes only when embedded within large sinking aggregates or fecal pellets (3–5). Phytoplankton cells embedded within bulk-collected sinking particles typically account for between 3% and 20% of total sinking particulate organic carbon (POC), and their presence can serve as an ecological indicator of the surface ocean processes that influence carbon export (6–8).

It is also possible for phytoplankton cells to sink as individuals or chains, but very few studies have provided evidence that this might be an important process of the biological carbon pump (9–12). Settling rates of phytoplankton cells have been measured extensively in laboratory studies that seek to quantify how changes in physiology, size, and mineral ballast affect the sinking speeds of individual cells (13–16). The relevance of solitary sinking cells within our conceptual models of the biological carbon pump is unknown, mostly because the flux of these individual cells is very difficult to measure in the ocean. In theory, small cells should sink too slowly to reach mesopelagic depths before being remineralized (17,18), and their fluxes would rapidly attenuate below the upper mixed layer (19). Presumably, any POC flux by individual cells would be relatively small compared to detrital particles, making solitary sinking cells essentially negligible in our accounting of the biological carbon pump.

Over the last two decades, small (<100 µm) particles have been increasingly recognized to play an important role in mesopelagic carbon fluxes, and in many ways phytoplankton cells fall within this often overlooked small-particle category (20–24). It is often assumed that small particles and cells sink too slowly to measurably influence bulk POC fluxes, but in some ecosystems particles sinking slower than ∼10 m d^-1^ can constitute the majority of particulate organic carbon (POC) flux into the mesopelagic (21,25). When resolved by size, small particles (<100 µm) often account for between 5-40% of the total POC flux (26,27), and sometimes the majority (28). It therefore seems plausible that individual solitary cells, among these small particles, could play a distinct and important role in POC export.

Solitary phytoplankton cells may be transported into the mesopelagic by some of the same export pathways that transport other small particles. For example, mixed-layer shoaling can isolate particle-rich surface water at depth, preventing cells from mixing back upward (29). Eddies can transport surface POC downward along tilted isopycnals (30). Intact phytoplankton cells can be packaged into larger, faster-sinking particles or fecal pellets, which may degrade and disaggregate at depth (31), releasing cells and small particle fragments into the mesopelagic. In this way, phytoplankton cells may be found in the mesopelagic due to one or more cycles of particle aggregation, disaggregation, and physical advection, in combination with direct gravitational settling.

Because they are living organisms, phytoplankton may also be exported into the mesopelagic by processes that are distinct from other small particles. Phytoplankton actively adjust their buoyancy or vertically migrate to access nutrient pools (10,32). When cells form resting or sexual stages they become denser and actively sink out of the surface ocean (33). Dead or dying cells lose their ability to regulate buoyancy, which also causes them to sink faster (14,34). Although the average sinking speed of phytoplankton cells is typically only a few meters per day (13), individuals within the population have a broad range of sinking speeds, some of which may be fast enough to reach mesopelagic depths (15).

The importance of solitary sinking phytoplankton cells in plankton ecology and ocean biogeochemistry remains uncertain primarily because this process is difficult to directly observe and is rarely disentangled from fluxes driven by detrital particles. We designed 42 sediment trap deployments capable of observing and quantifying this process in the subarctic North Pacific and North Atlantic Ocean. Throughout these month-long field expeditions, we deployed mesopelagic sediment traps that contained a gel layer in the bottom of the collection tube. Sinking particles embedded within this gel layer were collected intact and remained separated from one another, enabling us to distinguish between the flux of solitary phytoplankton cells and those embedded within detrital particles. We use our observations to assess the potential mechanisms delivering individual cells into the mesopelagic and to contextualize how these individually sinking cells fit into our framework of the biological carbon pump.

## Methods

### Study location

Sediment traps were deployed in the subarctic North Pacific (Ocean Station Papa; 50°N, 145°W; Aug-Sept 2018) and North Atlantic Ocean (Porcupine Abyssal Plain; 49.01°N,14.87°W; May 2021) during the EXport Processes in the Ocean from RemoTe Sensing (EXPORTS) field campaigns (35,36). In the North Pacific, phytoplankton growth was dominated by small cells (37) and tightly matched by microzooplankton grazing (38), resulting in a recycling food web with low POC export and efficiency (39,40). Export was punctuated by patchy salp swarms that produced episodic fluxes of rapidly sinking fecal pellets (41). In the North Atlantic, the demise of a spring diatom bloom resulted in high POC flux out of the surface composed of large phyto-detrital aggregates (42).

### Sediment trap deployments

Sinking particles were collected for periods ranging 1–6 days, at up to 5 depths in surface-tethered traps (STTs)(43) and neutrally buoyant sediment traps (NBSTs) (44) deployed 3 times at each study location (S1 Table). The sediment trap at each depth carried 4 cylindrical collection tubes with area 0.0113 m^2^; one contained a jar with a polyacrylamide gel layer for microscopic imaging and quantification, two contained a formalin brine layer for measurement of bulk chemical fluxes, and the last collection tube contained either formalin brine or RNA-later preservative to sample DNA within sinking particles (45).

### Bulk carbon fluxes

Bulk POC flux measurements are reported in detail elsewhere (40,46,47) and we summarize those analyses here. Upon trap recovery, overlying seawater within each trap collection tube was siphoned off. Duplicate tubes containing formalin brine (70 ppt salinity, 0.1% formaldehyde, 3 mM borate) were combined and drained through a 350 µm mesh to screen out zooplankton that actively swam into the trap tube. Screens were examined by microscopy, zooplankton were removed using forceps, and the remaining particles were rinsed back into the particle sample, then split into 8 equal fractions by volume (48). Three particle fractions were collected onto combusted glass fiber filters (Whatman GF/F) and total particulate carbon was measured by combustion elemental analysis. Three particle fractions were collected onto polycarbonate filters and used to measure particulate inorganic carbon (PIC), assessed by coulometry. POC was calculated by subtracting PIC from the total particulate carbon measurement. Fluxes were calculated by dividing by the split fraction, dividing by the collection duration (see S1 Table), and dividing by the total collection area (0.0226 m^2^). Uncertainties were propagated from the standard deviation of replicate splits.

### Image analysis and modeling POC fluxes by particle class

Particulate organic carbon fluxes by distinct classes of detrital particles were measured by quantitative image analysis of gel layers deployed inside of sediment trap tubes. These analyses and data are reported elsewhere (27,49), and we provide a summary here. After recovery from the sediment trap, the jar containing a polyacrylamide gel was imaged on a stereomicroscope (Olympus SZX16 with Lumenera Infinity 2 camera attachment) under multiple magnifications with fields of view evenly distributed across the gel surface. For samples collected in the North Pacific Ocean, gels were imaged at 7x, 20x, 50x, and 115x. For samples collected in the North Atlantic Ocean, gels were imaged at 7x, 32x, and 115x. This combination of magnifications enabled quantitative detection of particle sizes as small as ∼20 µm and including particles >1000 µm. Particles were detected in images using python-based functions in the scikit-image package, previously reported in detail (27). These included a combination of thresholding and edge detection steps to identify particles. Pixel regions containing potential particles were individually saved as separate particle images and used for classification. Tens of thousands of particle images were classified through a combination of manual labeling and assisted by iterative predictions by machine learning models as the modeling approach was being developed (49). All assigned particle classes were manually validated.

The contribution of each particle class to POC flux was calculated, and is described in detail elsewhere (27,49). Briefly, the volume of each particle type was calculated by estimating size from the pixel area and shape equations parameterized for each particle type. Particle volumes were converted into carbon units using class-specific shape equations and parameterizations. Carbon fluxes were calculated for each particle type by dividing the carbon in each particle by the imaged surface area of the gel layer, then dividing by the trap deployment time, and summing all the particles of a specific class or size.

### Phytoplankton cell fluxes

Sinking phytoplankton cells were counted in polyacrylamide gel layers using light microscopy and identified as diatom, dinoflagellate, coccolithophore, or silicoflagellate and additionally by genus or morphological shape when genus-level ID was not possible. Only distinct, individual cells or chains were counted; any cells adjacent to or incorporated within a detrital particle were not counted.

Cells were counted using a stereomicroscope under dark field illumination (SZX16, Olympus, Tokyo, Japan). The smallest resolvable cells (>10 µm) were quantified in a subsample of the gel at high magnification (115x). Cells were counted in at least twenty, 0.025 mm by 0.025 mm grid squares distributed randomly across the entire gel surface. We count at least 100 cells in total per sample, except 1 of the 13 North Atlantic samples and 7 of the 30 North Pacific samples. The entire gel surface was then surveyed at 25x and 7x magnifications to detect large, rare cells. The average number of cells counted per gel was 657±351 in the North Atlantic and 301±350 in the North Pacific, resulting in low counting uncertainties (square-root of the count number), which propagated into low flux uncertainties. Cell fluxes were calculated by dividing cell counts by the gel surface area surveyed and the trap collection period. Differences in phytoplankton community composition were assessed using permutational analysis of variance (PERMANOVA) of the Bray Curtis dissimilarity metric using tools in Python *(sklearn.neighbors.DistanceMetric, skbio.stats.distance.permanova).* These differences were visually assessed using a nonmetric multidimentional scaling analysis (*sklearn.manifold.MDS*). Phytoplankton number fluxes were related to other biogeochemical fluxes (particles, POC) throughout the mesopelagic for each deployment. Covariability among measures in each sample was resolved using a principal component analysis (*scipy.linalg.eig*) after first normalizing each variable to values between 0 and 1.

Cell diameters were measured from a subset of the most abundant cell types (∼10-80 µm) and the midpoint (45 µm) was used to calculate the spherical volume for estimating cellular POC (50) and the relative contribution of cells to total POC fluxes.

### Composition of phytoplankton in gel layers compared to bulk-collected particles

To test whether solitary cells detected in gel layers reflect the process of gravitational settling by single cells, the phytoplankton cell contents within bulk-collected sediment trap particles were compared to those collected side-by-side in a co-deployed gel layer.

Unpreserved, bulk particles were collected for microscopy in the North Atlantic during the third surface-tethered trap deployment from 500 m deep. In preparation, seawater was collected from 500 m using Niskin bottles on the CTD rosette, filtered through a 0.2 µm Supor membrane, and salinity adjusted to 45. This filtered seawater formed the bottom layer of 4 additional collection tubes attached to the STT drifting array at 500 m. The trap array was recovered after 1.8 days and particles from replicate tubes were combined and concentrated to 200 mL. A subsample of 30 mL was diluted up to 250 mL with 500 m water before being preserved in 1% glutaraldehyde. Preserved particles were settled for at least 24 hrs and concentrated by removing overlying liquid. At total of 269 phytoplankton cells were identified and counted using a Sedgewick Rafter on a Nikon Eclipse Ts2 microscope at 400x magnification. In contrast to microscopy counts of gel trap samples, no distinction was made between cells found inside or outside detrital particles.

### Data and code availability

Data are available in the NASA SeaBASS data repository at doi: 10.5067/SeaBASS/EXPORTS/DATA001. A static repository of the python code and the required data input used for this analysis will be published on Zenodo upon acceptance for publication.

## Results

### Distinct phytoplankton taxa sink as solitary cells

We interpret the solitary cells detected in gel layers as the distinct subset of phytoplankton that were sinking as individuals, whereas bulk-collected particles also include cells that were embedded within sinking detritus. We assume that the solitary cells arrived in the gel layer while sinking as distinct individuals, and not due to random disaggregation of detrital particle contents upon contact with the gel layer. To test these assumptions, we compared the composition of cells within a gel layer to those collected concurrently in a bulk-collected sample (North Atlantic STT deployment 3, 500 m trap depth). Solitary cells detected in the gel layer accounted for 22% of the total cell fluxes detected in the bulk sample (Fig. 1). The composition of these solitary sinking cells differed from the total community collected in the bulk sample. The community of solitary cells detected in the gel was dominated by the diatom *Thalassionema.* The flux of *Thalassionema* cells detected in the bulk-collected samples was equal to the solitary cell fluxes detected in the gels, indicating that the entire sinking population of *Thalassionema* was sinking as solitary individuals rather than embedded within detrital particles. Other phytoplankton groups (unidentified pennate diatom, *Chaetoceros*, *Bacteriastrum*, *Dichtyocha*, *Thalassiosira*, *Pseudo-nitzschia*) were less abundant in the gel layer but composed a large proportion of the cells fluxes in bulk-collected samples, indicating that they were primarily delivered to the mesopelagic embedded within detrital particles. Our subsequent reporting of cell fluxes detected in gel layers are therefore interpreted as solitary sinking cells, distinct from the community of cells embedded and transported to the mesopelagic within detrital particles.

**Figure 1.**
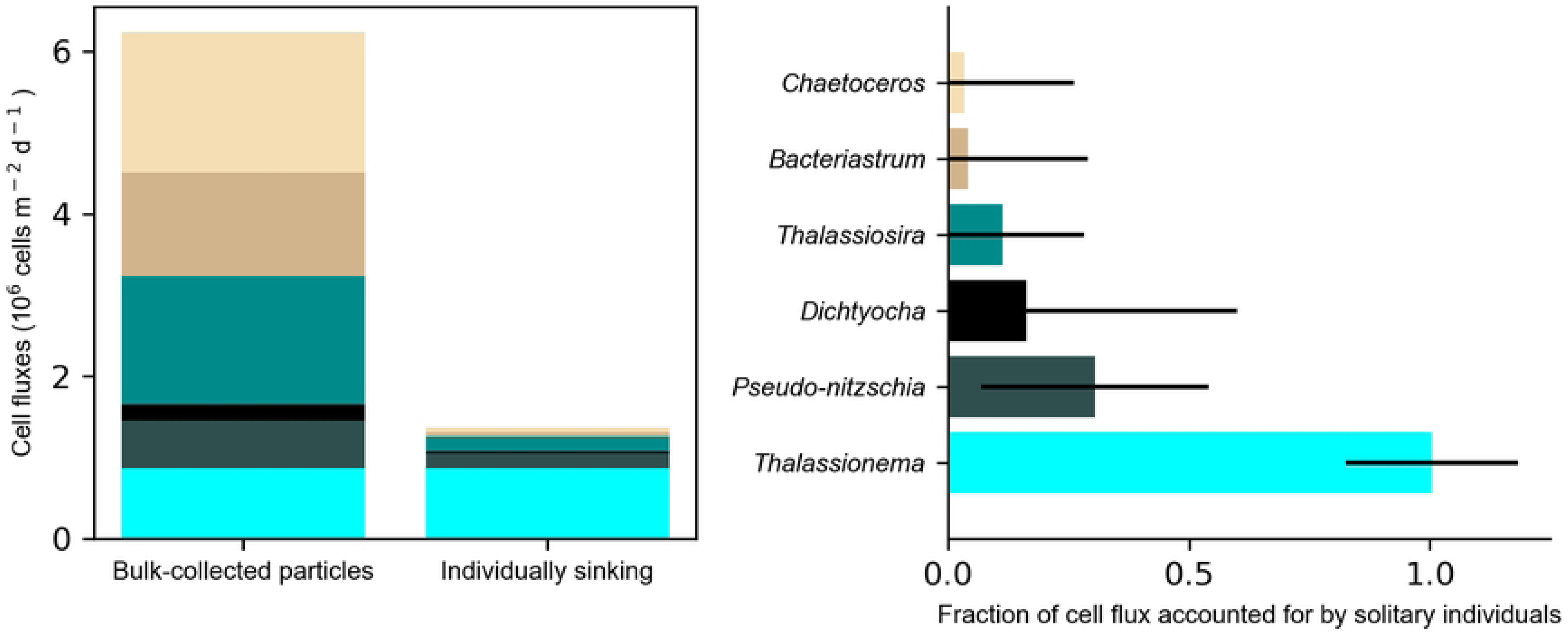
Comparison of phytoplankton cells fluxes detected in a bulk collected sample compared to solitary individuals counted in a corresponding gel layer. Left) Cell fluxes in a bulk-collected sample (North Atlantic, 500 m depth) include cells that were packaged within detrital particles and those sinking as individuals. Individual phytoplankton counted in gel traps are interpreted as solitary sinking cells. Genera or categories detected with <50% counting uncertainty are shown here. Right) Fraction of cell fluxes in a bulk trap also detected as individuals in the gel trap. A ratio of 1 indicates that all of the cell fluxes can be accounted for by solitary sinking individuals. Error bars represent the propagation of counting uncertainty.

### Solitary phytoplankton cell fluxes vary over depth and time

Solitary cell fluxes were an order of magnitude higher in the North Atlantic compared to the North Pacific (Fig. 2). Variability in cell fluxes collected by trap platforms deployed at the same time and depth, and up to 2 km apart, was similar to variability of detrital particle fluxes detected by previous studies (27,51); standard deviations of cell fluxes sampled by triplicate trap platforms were 22-79% of the mean. In the North Atlantic, phytoplankton cell fluxes decreased with depth and depth-averaged cell fluxes increased 3-fold from the first (0.42 x 10^6^ cells m^-2^ d^-1^) to the second (1.26 x 10^6^ cells m^-2^ d^-1^) deployment, and an additional 1.8 fold during the third deployment (2.23 x 10^6^ cells m^-2^ d^-1^). In the North Pacific, phytoplankton cell fluxes increased with depth (Fig. 1B). Depth averaged fluxes increased 4.2 fold from deployment one (0.055 x 10^6^ cells m^-2^ d^-1^) to deployment two (0.23 x 10^6^ cells m^-2^ d^-1^) and then decreased during deployment three (0.16 x 10^6^ cells m^-2^ d^-1^).

**Figure 2.**
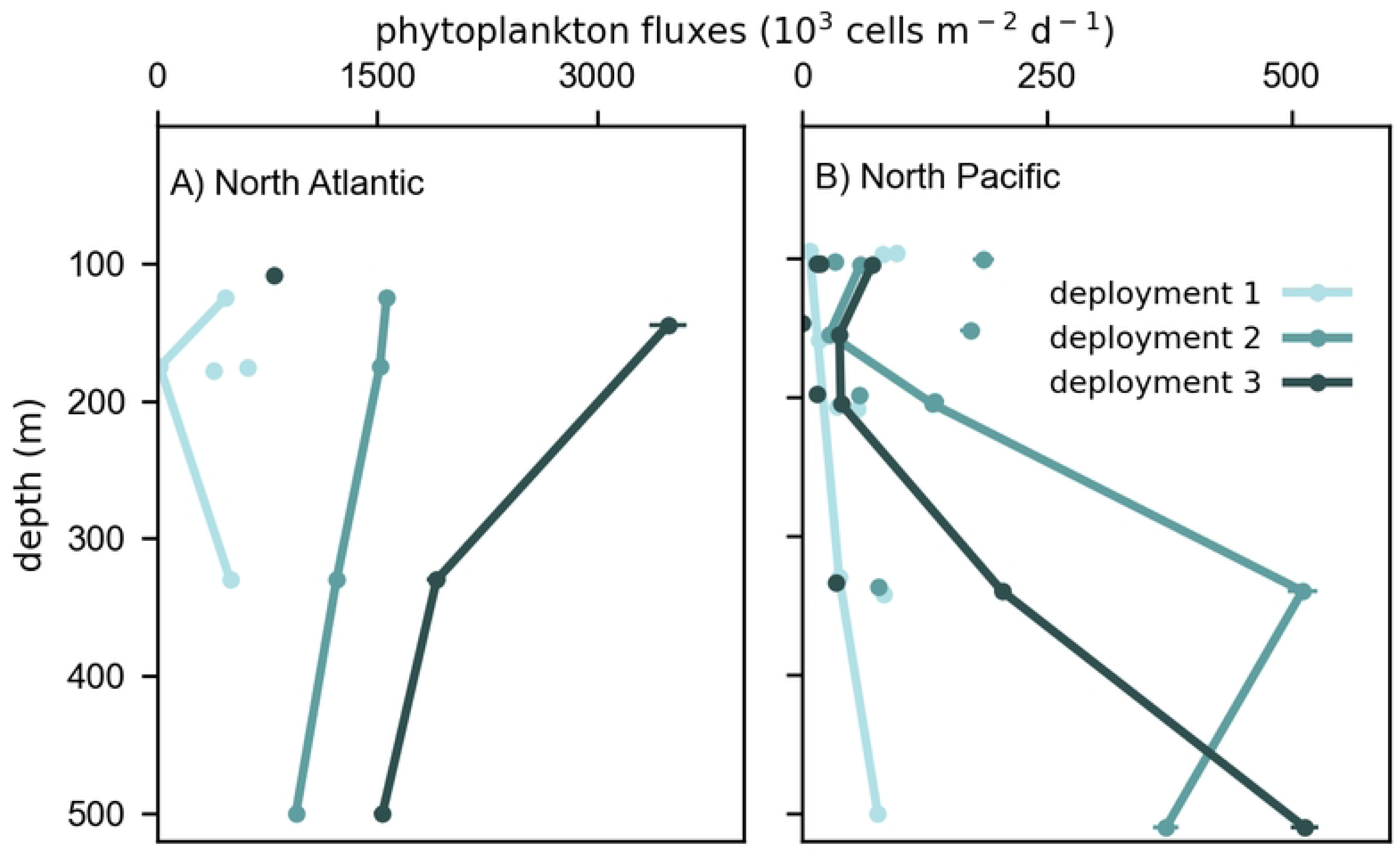
Variability in the number flux of solitary phytoplankton cells. Fluxes were quantified in 42 sediment trap gel layers in the North Atlantic (left) and North Pacific (right). Lines connect samples from the same surface tethered sediment trap (STT) array during each deployment period. Unconnected circles are number fluxes measured in neutrally buoyant sediment traps (NBSTs) deployed individually at similar depth as the STTs during each deployment. Not all depths have replicate trap samples. Error bars indicate uncertainties from cell counts, many of which are smaller than the size of the circle due to the large number of cells counted (see Methods).

### Solitary cell fluxes can be decoupled from detrital particle fluxes

Solitary sinking cells may be delivered into the mesopelagic by similar mechanisms as small detrital particles, causing fluxes of both particles types to vary over space and time. Alternatively, solitary cells and detritus could have distinct export mechanisms. If those export mechanisms are driven by the same ecosystem forcings, the fluxes of small detrital particles and cells could still be correlated. If their export mechanisms are not linked in any way, we expect their fluxes to be uncorrelated. To examine the relationship between solitary cell fluxes and detrital POC fluxes, we first estimated POC flux of solitary cells, assuming an average cell diameter of 45 µm. The calculated POC flux by solitary cells was less than 5% of the total POC flux measured in bulk sediment traps at most depths in both the North Atlantic (mean=3% ± 1.9%) and North Pacific (mean=1.6% ± 2.2%)(S1 Fig.).

Next, we compared the POC fluxes by solitary cells to the POC fluxes attributed to small (<100 µm) detrital particles, determined previously by image-based analysis of the gel layer (27,49). In the North Atlantic, POC fluxes by solitary cells were positively correlated with POC flux by small detrital particles (Spearman rank = 0.7, p=0.01, Fig. 3). In this ecosystem, solitary cell and small detrital particle fluxes were also correlated with large particle POC flux (not shown), suggesting that all three distinct particle pools had shared flux mechanisms or were coupled by larger ecosystem processes. In contrast, POC fluxes by solitary cells in the North Pacific were not correlated with small detrital POC flux (Spearman rank = −0.3, p=0.1, Fig. 3). Neither solitary cell nor small detrital POC flux was correlated with large particle POC flux (not shown), suggesting that these three particle pools were driven by distinct flux mechanisms with different time and space scales of variation.

**Figure 3.**
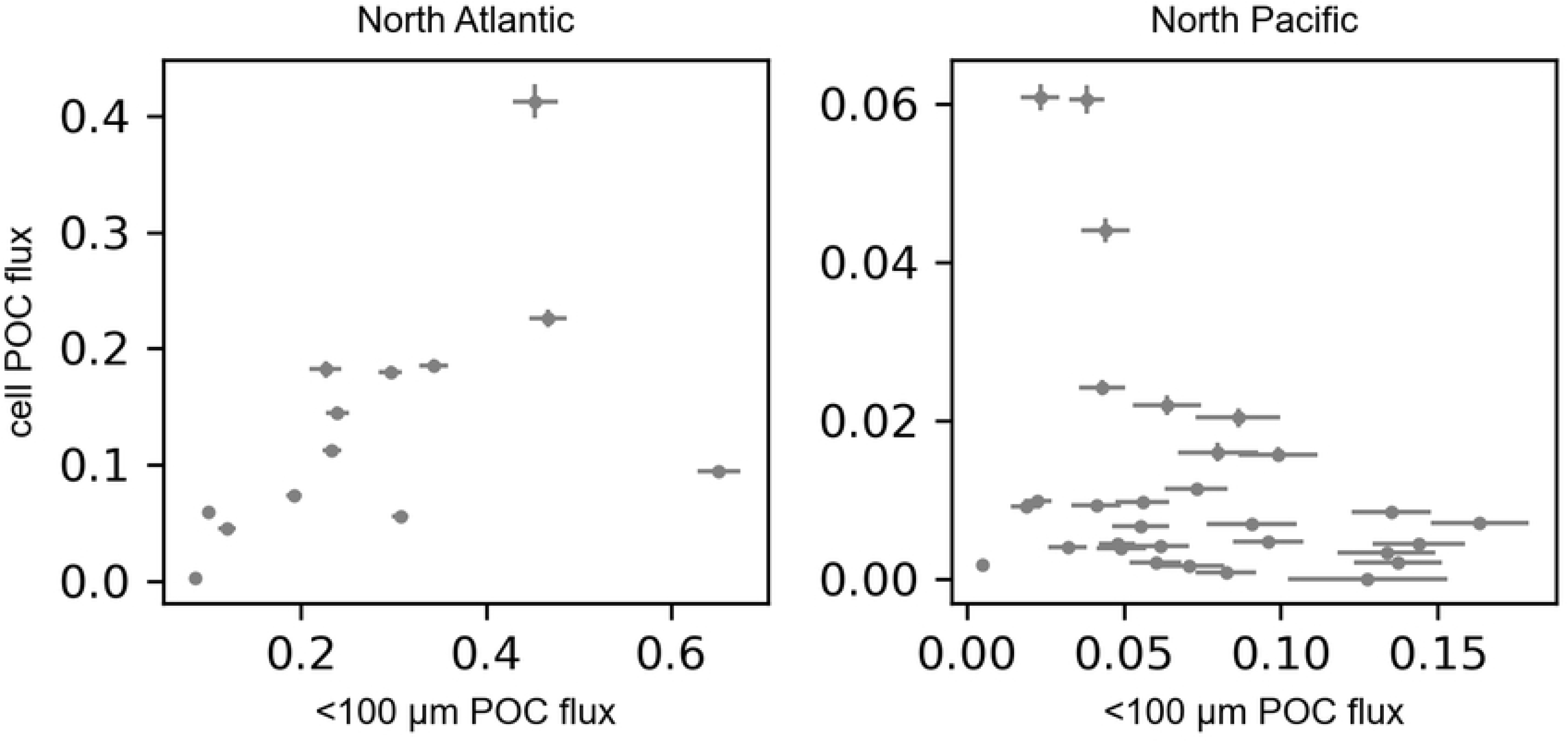
Relationship between phytoplankton cell POC fluxes and small (<100 µm) detrital particle POC fluxes in the North Atlantic (left, Spearman rank = 0.7, p=0.01) and North Pacific (right, Spearman rank = −0.3, p=0.1) Ocean. POC fluxes are in units mmol C m^-2^ d^-1^. Phytoplankton cell POC fluxes were calculated assuming an average diameter of 45 µm and using diatom-specific equations (50).

To further interrogate the association of solitary phytoplankton cell fluxes with sinking detritus, we tested for correlations of cell fluxes with POC fluxes of specific detrital particle types modeled from image-based analyses (27,49). In the North Atlantic, the number flux of solitary cells co-varied with POC flux of aggregates and long fecal pellets (Fig. 4), the particle types that comprised the majority of the POC flux (S2 Fig.). In the North Pacific, cell fluxes co-varied with short ovoid fecal pellets, which are typically produced by deep-dwelling zooplankton such as appendicularians (52,53). Short fecal pellets made up a small fraction of the total POC flux but were the only particle type whose fluxes increased slightly with depth (27). In both oceans, cell fluxes were negatively associated with the flux of salp fecal pellets.

**Figure 4.**
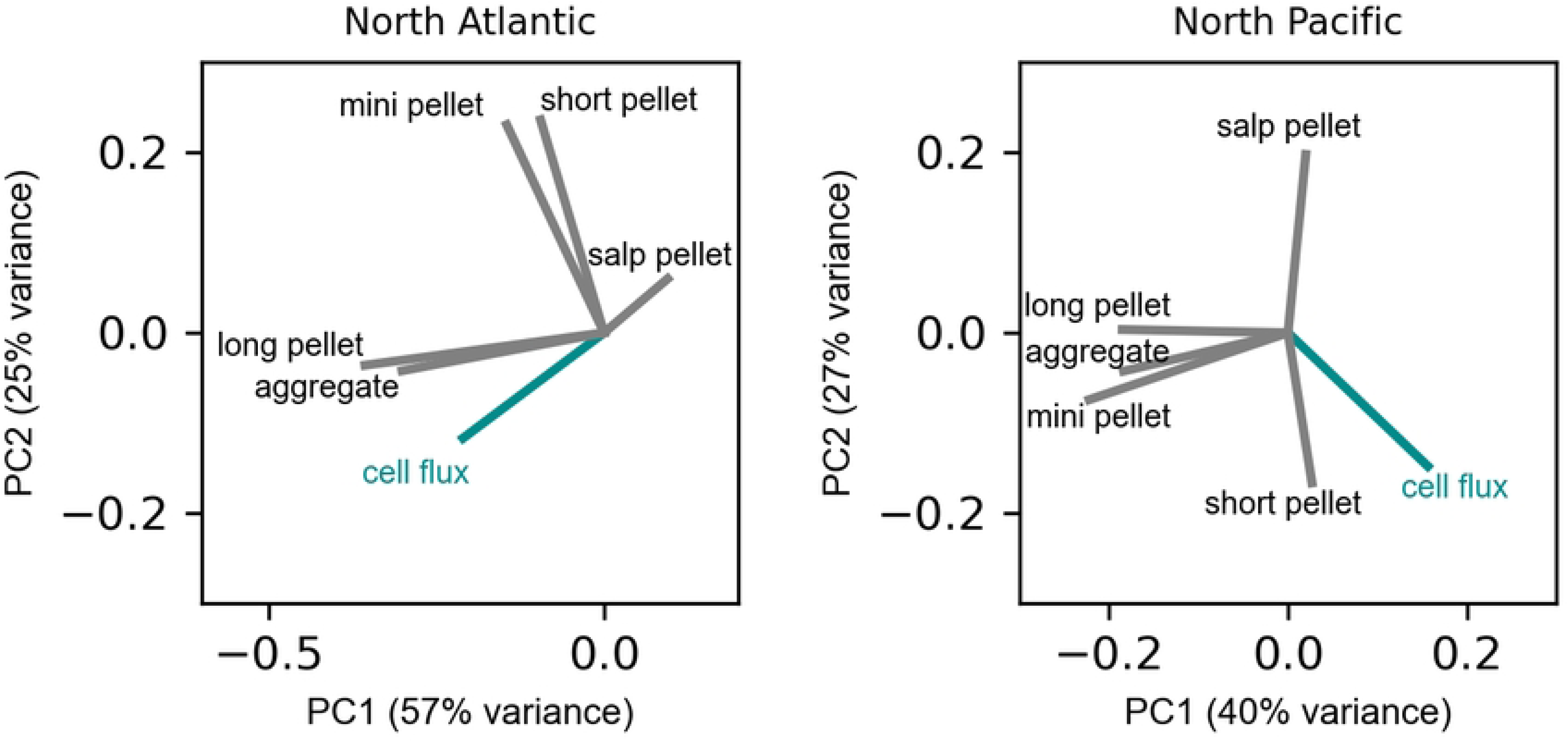
Covariance of phytoplankton cell number fluxes with POC flux by specific particle types in the North Atlantic (left) and North Pacific (right) Ocean. Principal component analysis of all measured fluxes shows the loading magnitude of each flux on the first two component axes.

### Variability in the composition of solitary sinking cells

Diatoms dominated solitary phytoplankton cell fluxes in both ocean ecosystems in the North Atlantic (93%) and the North Pacific (83%) when averaged over all samples. The remaining fraction consisted of coccolithophores, silicoflagellates, and dinoflagellates (Fig. 5). Among diatoms, *Thalassiosira* cells were a consistently large contributor to cell fluxes in both basins, particularly during the first trap deployments. In the North Atlantic, the composition of sinking phytoplankton changed over time more than with depth (S3 Fig., Bray Curtis dissimilarities, PERMANOVA p<0.05). After the first trap deployment, the sinking community was largely composed of *Thalassionema* diatoms.

**Figure 5.**
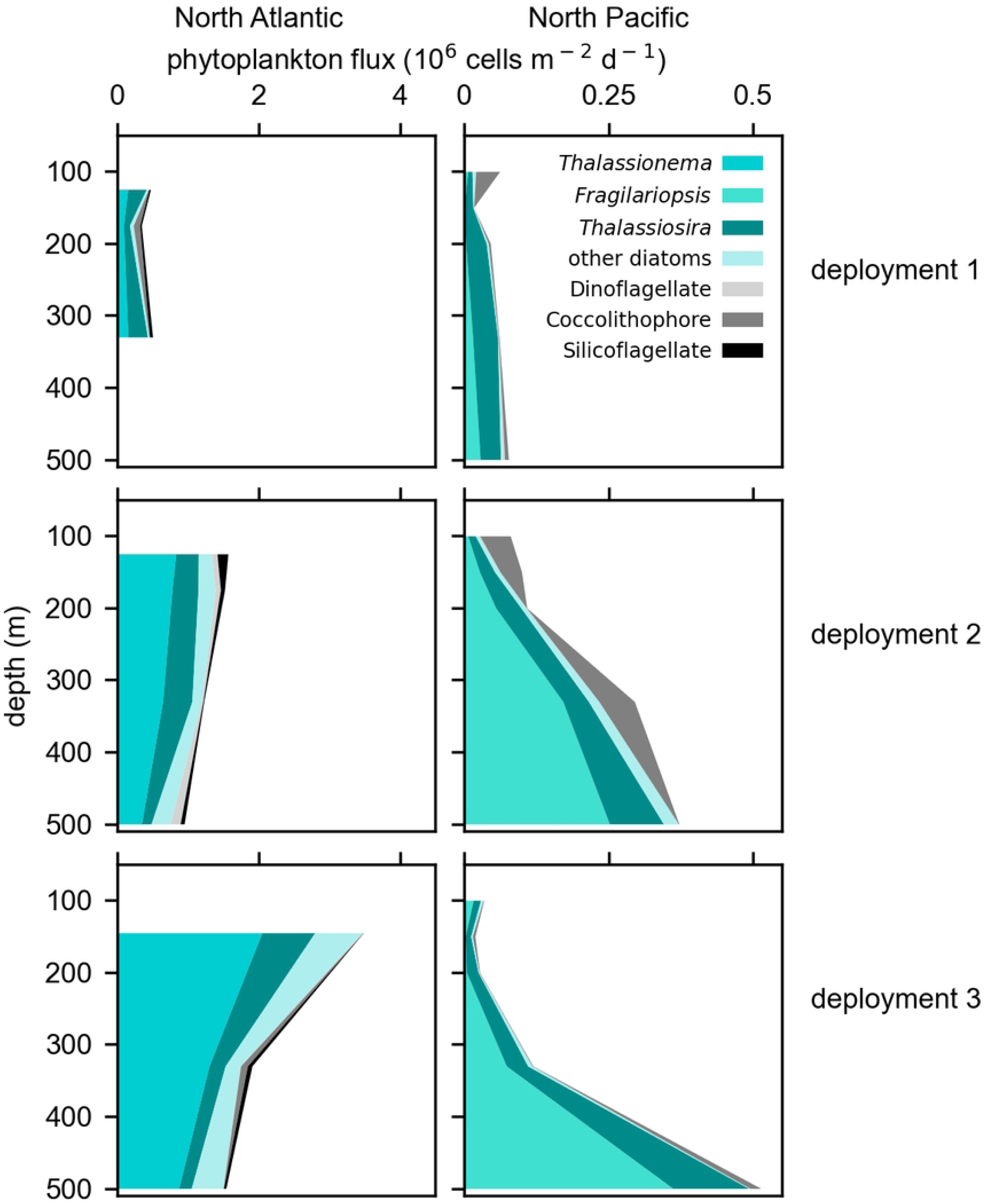
Number flux of solitary phytoplankton cells in the North Atlantic (left) and North Pacific (right) Ocean during three deployment periods. Cell fluxes from the North Pacific depict the average of STT and NBST trap samples deployed at the same nominal depth horizon during the same time period. In the North Atlantic, very few NBST gel trap samples were collected and depth profiles of phytoplankton cell fluxes are depicted by STT gel trap samples alone. The most abundant diatom genera contributing to number fluxes are distinguished from other grouped categories of phytoplankton.

In the North Pacific, sinking phytoplankton communities changed with depth more than over time (Fig. 5, S3 Fig.). The unusual flux profile of phytoplankton cells corresponded with a change in community composition in the deep community (>100 m) compared to the community collected in the shallowest sediment traps (Bray-Curtis dissimilarities, PERMANOVA p<0.05), driven largely by the increasing flux of *Fragilariopsis* diatoms.

To better visualize how the community composition of individually sinking cells changed over depth and with time, we created a network linking samples with the most similar communities of sinking cells (Bray-Curtis dissimilarity < 0.25) (Fig. 6). In the North Atlantic, a distinct community enriched in *Thalassionema* appeared during the second deployment, and persisted through the third deployment when it extended to deeper depths. In the North Pacific, 6 distinct networks of sample communities were detected. Sample cluster 1 included two replicate samples at 95 m during the first deployment, dominated by the flux of coccolithophores. All other community networks spanned across multiple depths and over time. Sample clusters 2 thought 5 linked various sample replicates across overlapping depth regions and across all three deployments. These overlapping networks contained *Thalassiosira* as a relatively large percentage of cell fluxes. Sample cluster 6 was enriched in *Fragilariopsis* and did not overlap in space or time with other clusters. This distinct sinking population appeared deeper than 300 m during the second deployment and only appeared in the deepest trap sample (500 m) during the third deployment.

**Figure 6.**
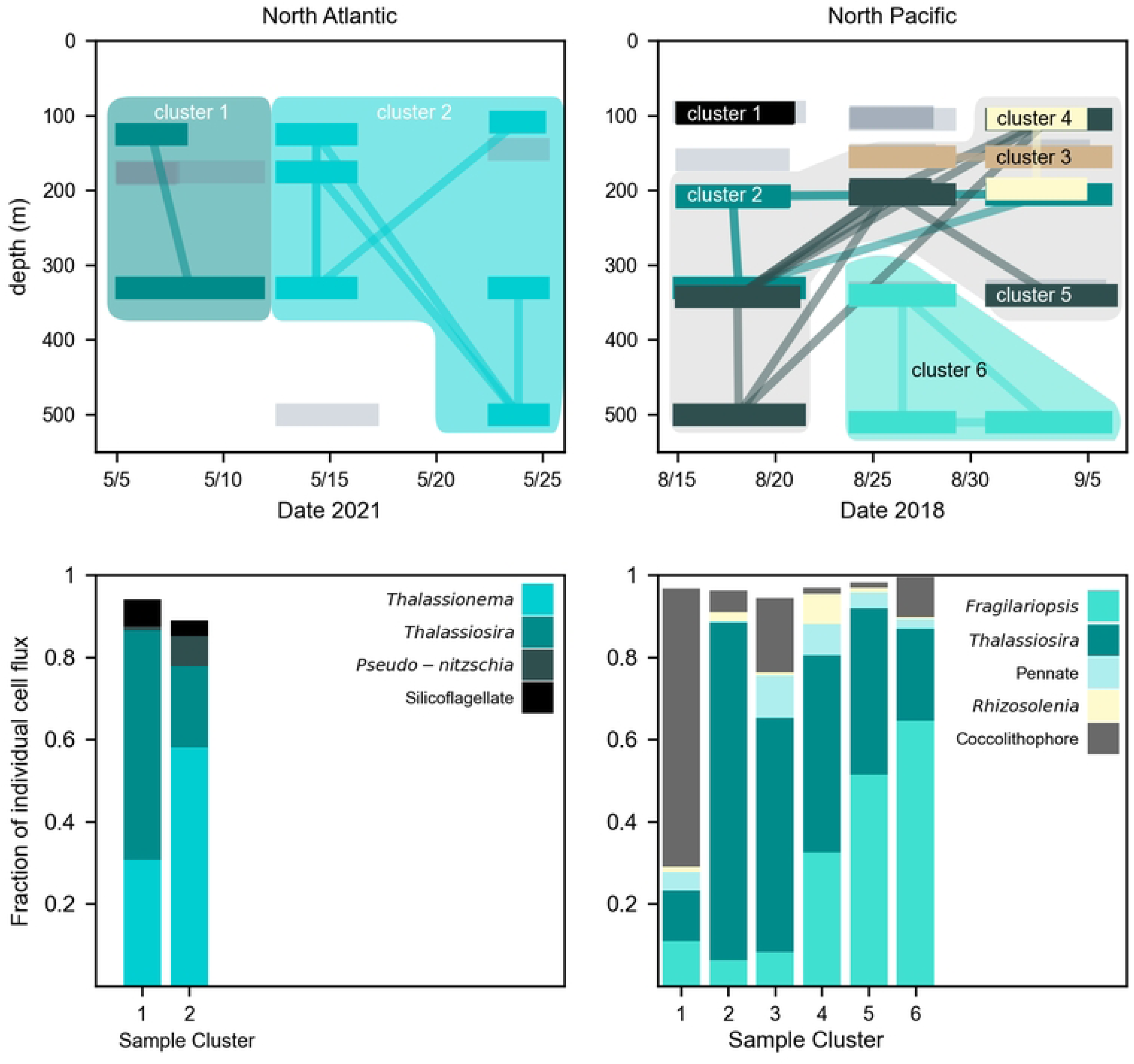
Network of solitary sinking phytoplankton communities over time and depth. Top row: Horizontal bars indicate trap deployment periods and depth while vectors link trap samples with similar phytoplankton communities (<0.25 dissimilarity score, Bray-Curtis). Grey bars without linkages indicates samples whose communities did not meet this similarity threshold with any other sample communities. Background color blocks illustrate regions of distinct community networks. In the North Pacific, samples clusters 2 through 5 overlapped in depth and time but contained different communities. Bottom row: Average community composition of all trap samples within a connected network. Taxa composing at least 5% of the flux within a network are shown.

### Gravitational settling may transport solitary cells through the mesopelagic

The downward shift in a distinct phytoplankton community in the North Pacific corresponded with a depth profile of cell fluxes that appeared to shift downward. This also corresponded to a loss in depth-averaged cell fluxes between the second and third deployments. As a thought exercise, we tested whether these changes might be attributable to the gravitational settling of *Fragilariopsis* at a reasonable sinking speed. The major assumptions of this exercise are that other loss processes to individually settling cells (e.g., spatial patchiness, grazing, remineralization) were minimal, thus the sinking speed estimated is conservatively large. We used the depth-integrated loss of solitary *Fragilariopsis* diatom cell fluxes between deployment 3 compared to deployment 2 to calculate the depth that cells would have to sink below the 500 m observational regions in the 7 days between deployments (see Supplemental Information and S4 Fig.). We calculated that a sinking speed of 6.6 m d^-1^ could account for the downward shift in the depth distribution of *Fragilariopsis* cell fluxes.

## Discussion

We found that solitary phytoplankton cells can sink through the water column in spite of their small size. Utilizing gel layers in sediment traps allowed us to identify the distinct subset of phytoplankton cells that were sinking as solitary individuals in the mesopelagic. While solitary settling cells are not typically incorporated into frameworks of the biological carbon pump, several other studies have also suggested that they may be ecologically important (9,11,12). The scale of our observations (42 sediment trap samples over 1-month periods in two ocean basins spanning 500 m depth) significantly bolsters the conclusions of these earlier studies. Here we assess what mechanisms may explain variability in cell fluxes and we contextualize their relative importance in our broader understanding of the biological carbon pump.

Our observations suggest several possible mechanisms that may be responsible for the delivery of solitary cells into the mesopelagic and their continued settling after delivery. In the North Atlantic, the delivery of phytoplankton cells could not be disentangled from the fluxes of small particles, or the fluxes of large particles like aggregates and fecal pellets. This correlation between cell fluxes and detrital particle fluxes may have been caused by the disaggregation of large particles, releasing small particles and cells into the water column. However, Romanelli et al. (42) observed that small (<100 µm) particles sinking during our field observations were distinct from the flux event of large sinking aggregates in the mesopelagic, and these small sinking particles were not generated by disaggregation in the mesopelagic. The small particles, enriched in biogenic silica, appeared to be generated in the surface mixed layer or immediately below, and continued to sink in the mesopelagic over the same observational period as our trap deployments. We suggest that the solitary sinking phytoplankton cells we observed in this study were among these small sinking detrital particles and likely the reason this small particle pool was enriched in biogenic silica. Similarly, our data suggest that the solitary cells were sinking as individuals and not delivered to the deep mesopelagic by disaggregation of larger sinking particles. Most of the cells detected in the gel layer (*Thalassionema*) did not appear to be embedded within the sinking detritus. Instead, bulk particles contained a different community composition. These observations suggest that most of the cells we detected as solitary sinking had not been recently disaggregated from larger particles. If this interpretation is correct, the correlation of cell fluxes with POC fluxes would have had to be caused by secondary physical or ecological processes that indirectly produced similar flux variability among particle pools.

In the North Pacific, we were surprised to observe an increase in cell fluxes with depth that were not correlated with small detrital particle or total POC fluxes. Phytoplankton cell fluxes only correlated with the fluxes of short fecal pellets produced by deep dwelling zooplankton (e.g., appendicularians), which also increased with depth. Our dataset does not allow us to disentangle whether these deep zooplankton communities played a role in enhancing cell fluxes, or another as-yet unidentified process drove the correlation. Stephens et al. (54) created a carbon budget for the time period of these field observations in the North Pacific, and concluded that mesopelagic communities were sustained by a large pool of slowly sinking particles that were probably exported from the surface ocean earlier in the year, prior to our research expedition. The deep, sinking phytoplankton community that we detected may have been part of this early summer export and decoupled in time from the overlying surface ecosystem. The downward progression of the maximum phytoplankton cell flux and associated community composition during the observational period could have been plausibly explained by settling of cells. To be captured in our sediment trap gel layers at the base of the collection tubes, these solitary cells must have been negatively buoyant, with a minimum sinking speed of 0.1-0.4 m d^-1^ (70 cm distance over 1.7-5.9 days). This is well within previously reported cell sinking rates (13). Our calculation suggested that the most abundant diatom in this deep community, *Fragilariopsis*, would have had to sink 6.6 m d^-1^, which is within the range of sinking speeds observed for diatoms within this size range (1.5 – 19 m d^-1^)(15). Alternatively, this apparent temporal variation with depth could have been caused by lateral advection. Still, Stephens et al. (54) did not find support for lateral advection as a source of deep organic carbon during this observational period.

Our observations establish that solitary phytoplankton cells can sink in the mesopelagic in spite of their small size, but the question remains: what is their ecological and biogeochemical significance? We estimated that these solitary cells typically contribute less than 5% to the total POC fluxes, perhaps a negligible omission in our accounting of ocean carbon budgets given the large uncertainties. Still, intact and potentially living phytoplankton cells would be of great nutritional value to mesopelagic consumers compared to previously digested detritus, which would give their relatively low POC fluxes an outsized influence on the mesopelagic ecosystem functioning. Small and slowly sinking particles in the mesopelagic have repeatedly been identified as a key food source to deep food webs that must be accounted for to close the mesopelagic carbon budget (54,55). If still alive, phytoplankton cells may serve as a particularly persistent and nutritious source of carbon to these ecosystems due to their physiological capacity to maintain active defenses against microbes and silica dissolution(56), and to survive for long periods in the dark (11,57).

The sinking of individual phytoplankton in the mesopelagic may also be an unaccounted-for component of phytoplankton ecology and evolution. Changes in buoyancy and sinking speed have long been hypothesized to be adaptations to changing surface conditions (58), enabling certain taxa to escape unfavorable surface conditions until deep physical mixing brings them back to the shallows to re-seed a bloom. Agusti et al.(11) found that 18% of the phytoplankton they collected from deep sea waters using specialized collectors were still alive. Cells may also sink as an adaptation when they form resting or sexual stages (59,60). Resting spores were a major driver of phytoplankton fluxes during previous observations of the North Atlantic spring bloom (33), though we rarely observed these cell morphologies during our study.

These observations provide robust support for the ubiquitous presence of solitary sinking phytoplankton cells in the mesopelagic ocean. While particle disaggregation may be responsible for the delivery of some of these cells, our data suggests that direct gravitational settling of phytoplankton cells is also an important mechanism of the biological carbon pump. Sinking phytoplankton cells play a functional role in mesopelagic ecosystems and carbon export that is distinct from the detrital particles typically accounted for in models of the biological carbon pump.

## Acknowledgements

This work was funded by the NASA Ocean Biology and Biogeochemistry program awards 80NSSC17K0662 and 80NSSC21K0015. CAD received additional support from the David and Lucile Packard Foundation. We thank the NASA EXPORTS science team, the captains and crew of the R/V Roger Revelle and RRS James Cook, and chief scientists Debbie Steinberg and Jason Graff. Samples were collected with additional support from the EXPORTs sediment trap team; Melissa Omand, Ken Buesseler, R. Patrick Kelly, Sean O’Neill, Alyson Santoro, and Nicola Paul. We also thank Heather McNair for assistance with sample collection and Melissa Omand, Christine Huffard, and Sebastian Sudek for comments on a draft of this manuscript.

---

**S1 Figure.**
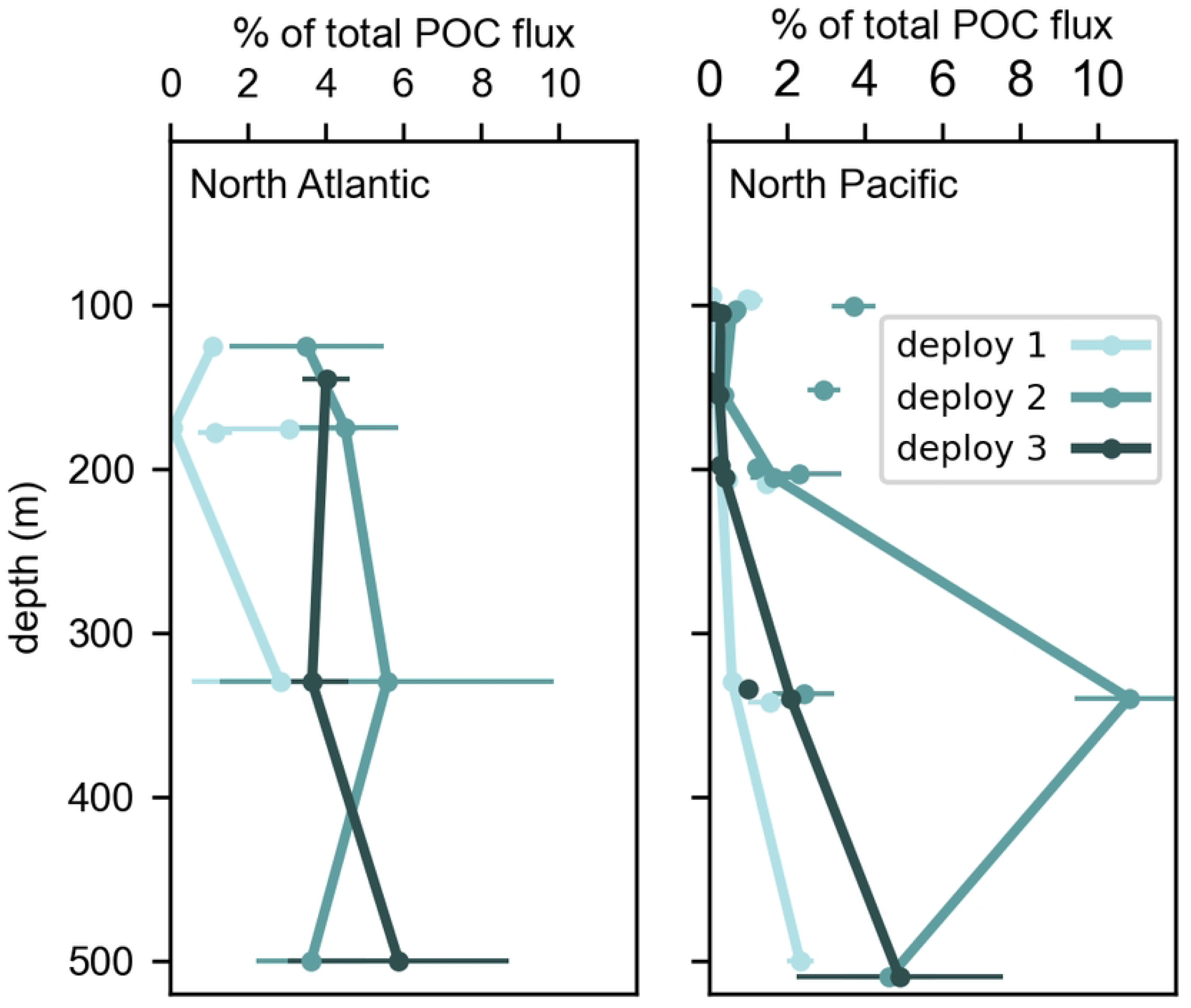
POC flux by solitary sinking cells as a percentage of the total POC flux measured in bulk sediment traps. Lines connect samples from the same surface tethered sediment trap (STT) array during each deployment period. Unconnected circles are fluxes measured in neutrally buoyant sediment traps (NBSTs) deployed individually at similar depth as the STTs during each deployment. POC fluxes by solitary cells were calculated assuming an average diameter of 45 µm and using the diatom-specific equations (50). Uncertainties represent the propagated uncertainty of both the cell counting uncertainty and the POC measurement uncertainty.

**S2 Figure.**
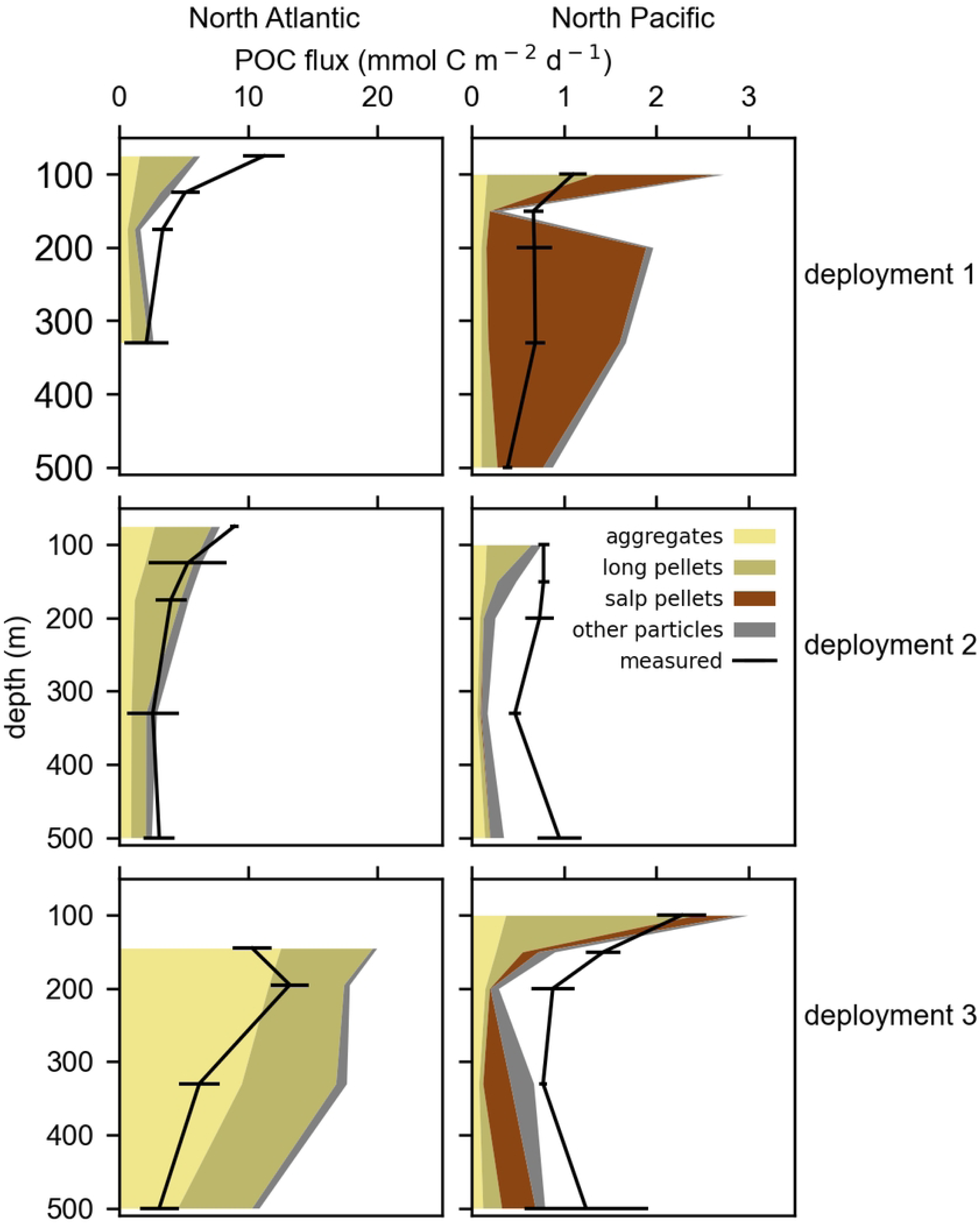
POC flux by detrital particles, calculated from imagery of gel layers in sediment traps and using modeled parameters(27,49). Chemical measurements of bulk POC flux are shown to ground truth fluxes calculated for specific particle types based on imagery. Uncertainties in the measured POC fluxes represent the standard deviation of replicate subsamples.

**S3 Figure.**
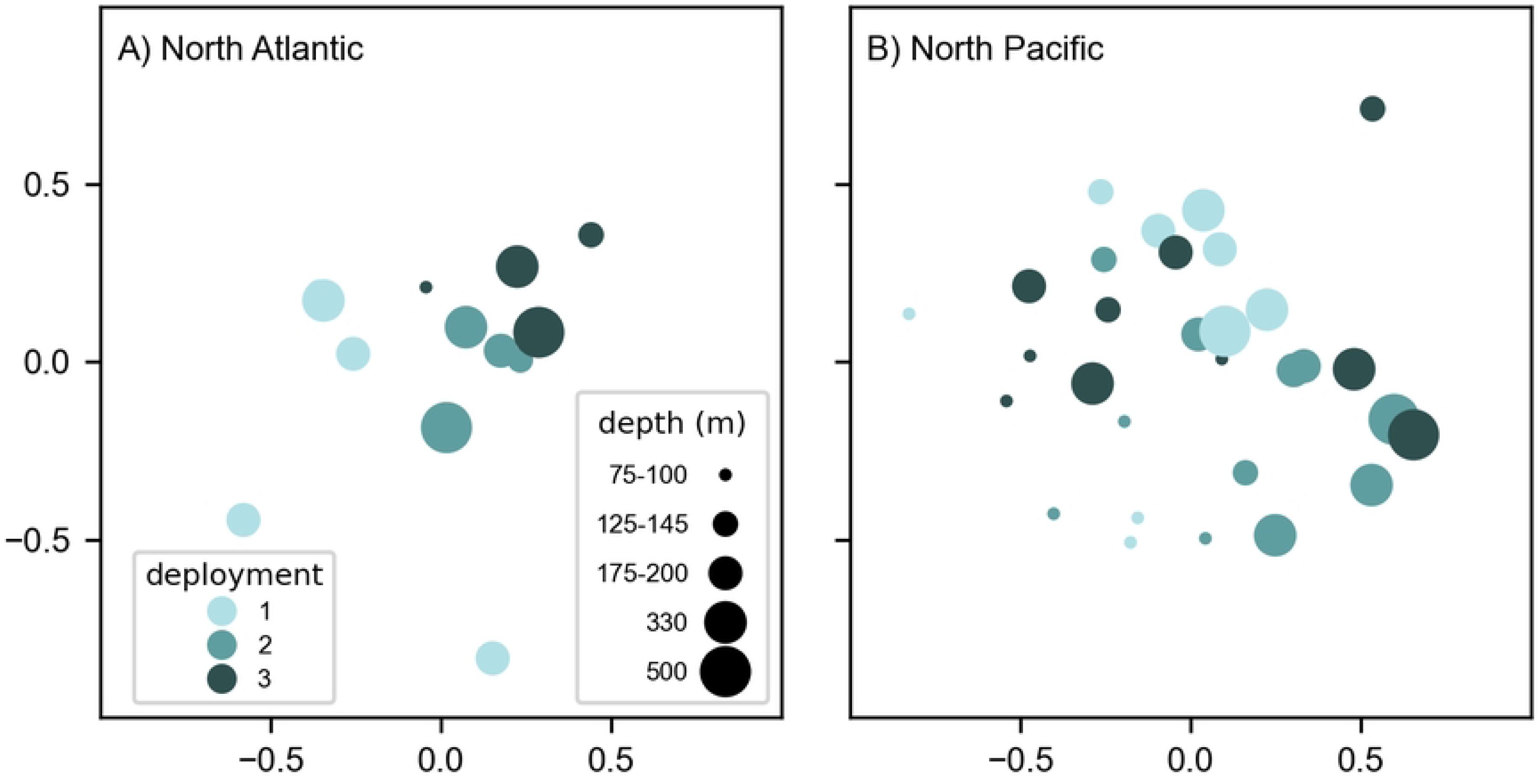
Differences in composition of individually sinking phytoplankton communities based on Bray Curtis dissimilarity. A) North Atlantic Ocean and B) North Pacific Ocean. Distance between samples (dots) on the Multidimensional Scaling plot represents dissimilarity among sample communities. Deployment periods are represented by color and depths are represented by dot size. Phytoplankton sinking in the North Atlantic differed most over time (deployment 1 vs. 2 & 3) while phytoplankton sinking in the North Pacific differed most across depth (95 m versus all other depths).

**S4 Figure.**
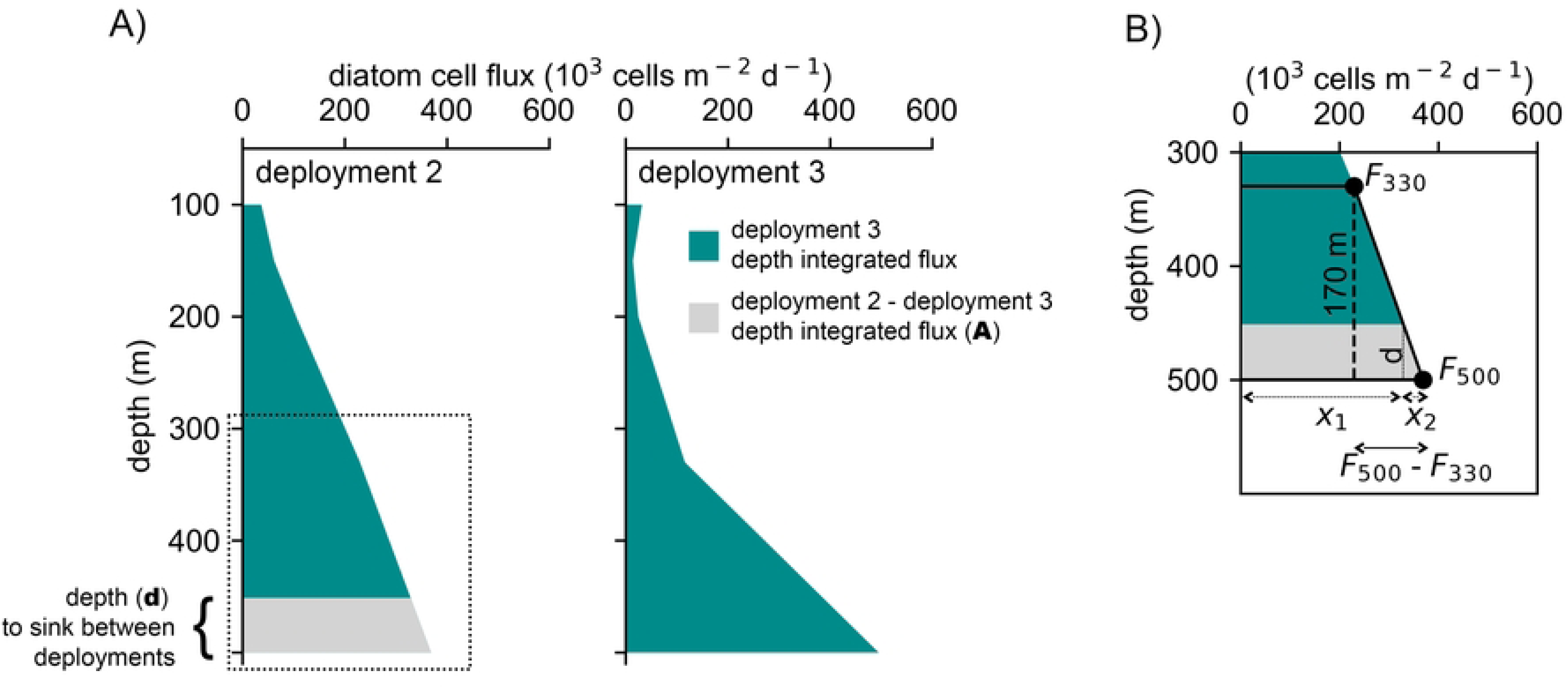
Changes in the depth-integrated *Fragilariopsis* cell fluxes used to calculate theoretical sinking speeds. A) The difference in depth-integrated cell fluxes between deployments 2 and 3 in the North Pacific. The gray integrated flux area illustrates the amount of depth-integrated flux reduced during deployment 3. Dashed box is shown in panel B. B) Known and unknown variables of the integrated depth flux profile used to calculate the depth (d) over which cells would have to sink in the 7 days between trap deployments.

**S1 Table. Summary of all sediment trap deployments in the North Atlantic and North Pacific study sites, including location, deployment days, trap platform type, depths of deployments in meters, and deployment duration in days.**

